# Intentions to use patient-initiated partner notification and acceptability of provider-initiated partner notification for Sexually Transmitted Infections – A cross-sectional survey among minibus taxi drivers in Gauteng Province, South Africa

**DOI:** 10.1101/502724

**Authors:** Mathildah Mpata Mokgatle, Sphiwe Madiba

## Abstract

**Background:** In South Africa, utilization of patient-initiated partner-notification (PN) using referral-slip in the management of sexually transmitted infections (STIs) is limited and only a limited number of sexual partners are ever notified. The study assessed the use of patient-initiated PN method using notification and referral slips and measured the level of acceptability of provider-initiated PN using short-message-service (SMS) to personal mobile phones of sexual partners.

**Methods:** A quantitative survey using anonymous structured self-administered and researcher assisted questionnaires was conducted among minibus taxi drivers in the nine major taxi ranks in Gauteng province, South Africa.

**Results:** The sample consisted of 722 minibus taxi drivers with a mean age of 37.2 years old, 284 (59.5%) had multiple sexual partners, 368 (52.2%) did not use a condom during last sexual act, 286 (42.8%) reported inconsistent use of condoms, and 459 (65%) tested for HIV in the past 12 months. Majority (n=709, 98.2%) understood the importance of PN once diagnosed with STI, but would prefer delivering PN referral slip (n=670, 93.2%) over telling a partner face to face if they themselves were diagnosed with STI. Acceptability of provider-initiated PN using SMS was 452 (62.7%) and associated with history of HIV testing in the past year (OR=1.72, p=0.002, CI: 1.21-2.45). The perceived use of PN referral-slip from sexual partner to seek treatment was 91.8% (n=659). About a third (n=234, 32.5%) were not in favor of provider-initiated PN by SMS and preferred telling partners face to face.

**Conclusion:** There were contrasting findings on the acceptability and utilization of existing patient-initiated PN and the proposed PN using SMS from health providers. The preference of delivering PN referral slip to sexual partner over face-to-face PN renders communicating about STIs the responsibility of health providers. Therefore, they have an opportunity to provide patients with options to choose a PN method that is best suited to their relationships and circumstances and modify PN messages to encourage partners to use the different PN to prevent STIs.

## Introduction

South Africa’s burden of disease due to sexually transmitted infections (STIs) is currently one of the largest in the world and this is true for all STIs, including HIV and human papilloma virus [1]. The significantly high prevalence of sexually transmitted infections (STIs) in Sub-Saharan Africa poses a threat because of the increased risk of HIV transmission [2]. In the African region, among the population group of 15 -49 year of age, STI prevalence and incidence for four curable STIs (chlamydia, gonorrhea, syphilis and trichomonas) was 19.4% and 24.1% [3-5]. The prevalence of syphilis among attendees of antenatal care in South Africa was 1.5% in 2011 [6]. From 2010-2011, the STI incidence for South Africa was 3.9% [7]. Kenyon and colleagues reported prevalence of syphilis and male urethral discharge in South Africa at 8.3% and 13.8% respectively [4].

Key drivers of STIs include risky sexual behavior such as multiple sexual partners, transactional sex, vulnerability of women in sexual relationships, and the high rate of concurrent partnerships [8-17]. Risky sexual behavior and the asymptomatic nature of STIs during early stages of the infection makes the control and treatment of STIs complicated [16]. Occasional lack of and/or shortage of partner notification (PN) and referral slips in the health facilities and the lack of strategies for healthcare workers (HCWs) to follow-up sexual partners further compromise STI control [14, 15].

The silent nature of STIs presents a huge public health threat, hence STI partner notification (PN) and referral is an important public health approach to reduce the burden [16]. Partner notification and referral is beneficial in controlling the spread of STIs if done correctly. Moreover, the use of electronic communication like SMS can facilitate partner notification [17] given the increased number of people globally who have access to mobile phones [18]. The current protocol for PN in the guidelines for the management and control of STIs in South Africa includes prescribing a syndromic management regimen, health education and STI/HIV counseling for the patient as well as patient-initiated PN using a notification and referral slip [11]. The process of PN using referral slips starts as the healthcare provider gathers the number of sexual partners from the patient and issues the relevant number of anonymous referral slips. The anonymous referral slips contains information about the risk of STI to the sexual partner and an invitation to receive treatment at a convenient health facility for sexual partners. The patients are then required to deliver the slips to their sexual partner within a period of a week and hence the name patient-initiated PN [11].

The utilization of patient-initiated PN is limited due to under-reporting of the number of sexual partners by the patients. Under-reporting the number of sexual partners reflects HCWs issuing less notification and referral slips [12, 13]. Under-reporting occurs due to reluctance to openly discuss sexual issues, the biological nature and characteristics of the STIs, and due to fear of moral judgment [1, 5]. Moreover, individuals rarely inform their sexual partners after diagnosis and treatment for STIs and in cases of multiple and concurrent sexual partnerships, the patient may not have the contact details of the casual sexual partner or they may not particularly care for the partner and hence see no need to notify them [12,15, 17–20]. Failure to inform sexual partners of their exposure to STIs increase the risk of STI transmission to other sexual partners who remain asymptomatic, and continuous infection of new partners and re-infections [19].

The extent of the STI problem and the association with HIV transmission highlights the need to continue and strengthen prevention and control of STIs [4]. The effective treatment and control of STIs depends on screening to detect and treat STIs among the sexual partners of the STI infected patients, which is dependent on the patient-initiated PN practices using referral slips in the South African treatment protocol. In addition to the patient-initiated PN practices, provider initiated PN such as text messaging, the internet, and phone calls are promising strategies to expand PN services [14, 15, 18]. The benefits of provider-initiated PN are that electronic messages may enhance rapid notification because they can reach partners who may be geographically dispersed, are likely to be used with partners who may not be notified otherwise, and come at a low cost [18]. Provider-initiated PN using short-message-service (SMS) presents as an additional and promising strategy for control and prevent STIs in South Africa [12, 13, 15].

Usually, sex workers and truck drivers are regarded as the highest risk for STI and HIV transmission and minibus taxi drivers are reported as another important vulnerable category [20, 21]. HIV infection amongst minibus taxi drivers is a concern, because of the occupational demand of being away from their families for long times and in some cases being exposed to unhealthy environment and falling prey to unhealthy sexual practices [21]. Minibus taxi drivers are identified as a high-risk population group with an increased risk of HIV infection because the minibus industry is characterized by a high level of migration, low level of education and stressful working conditions. Moreover, the focus of existing research has been on the assessment of HIV infection and AIDS in the transport industry amongst truck drivers, but not in the minibus industry in Gauteng province and in, South Africa generally [21]. Men in the transport industry have high sexual risk behaviors and low HIV risk perceptions, which have implications for STI transmission [21, 22]. Furthermore, minibus taxi drivers have poor access to health screening facilities because of the nature of their work, which may have an impact on STI treatment and PN [23]. Being highly mobile or travelling long distances impact on the ability of the patient to be able to reach the sexual partner for referral for STI treatment [21].

To add to the literature on sexual health among high sexual risk groups and the need to strengthen prevention and control of STIs, this paper presents an assessment of perceived use of patient-initiated PN using a referral slip from sexual partner, the acceptability of provider-initiated PN using SMS to personal mobile phones of the sexual partners and associated factors. The survey focus was on self-reported STIs among male minibus taxi drivers, who have access to the syndromic management of STI according to the South African treatment protocols and WHO treatment guidelines.

## Methods

### Study setting

The minibus taxi industry in South Africa is still largely unregulated even though it is the main form of public transport, due to limited options to affordable and accessible transport. With a fleet of at least 300,000 vehicles, the industry the taxi industry employs more than 600,000 people and transports 15 million commuters per day provides which is an equivalent of 65% of public transport commuters [24]. This makes the minibus taxis the most common transport especially because it is the only transport with large coverage of routes in the country. Minibus taxis are usually more reliable, have shorter waiting times, and longer running times, which are up to 24hrs especially for long distance and cross-border trips. Minibus taxi drivers run set routes as stipulated by the taxi association, which is the regulating body, but often pick up and drop off commuters anywhere in between. A typical layout of the industry consists of a series of taxi ranks, hubs of the greater system, where hundreds of taxis line up to transport people off to all parts of the city and beyond. Unlike metered taxis, minibus do not go door-to-door, but operate on routes, much like a bus. Big ranks can have dozens of taxi routes coming in and going out [25, 26]

Tshwane District in Gauteng province is one of the large cities that has a complex network of taxi ranks transporting passengers to and from the city, urban areas, peri-urban areas. Majority of the taxi ranks are situated near shopping malls, industrial areas, hospitals and business centers, for ease of access by passengers. In Tshwane district, only a few taxi ranks are built as formal structures to enable exchange or transfers and articulation with other modes of transport such as the bus or train. Minibus taxis from smaller taxi ranks in the townships, suburbs and informal settlements merge to feed central taxi ranks and passengers select their trips by choosing a relevant taxi to their destination [26].

### Design

A quantitative survey using anonymous structured self-administered and researcher assisted questionnaires was conducted. The survey was the first stage of a large formative evaluation project, which utilized a mixed method approach employing quantitative and qualitative methods to assess the acceptability and feasibility of implementing STI provider-initiated PN using SMS. High-risk populations were the target population for the study and they include minibus taxi drivers, out of school youth and young adults accessing primary health facilities, and university students in Tshwane District, Gauteng province, South Africa. This paper presents the findings frm a sample of the minibus taxi drivers.

### Study population and sample

The survey was conducted in nine major minibus taxi ranks and all drivers working in the selected ranks formed the study population. Systematic random sampling of taxis that were waiting to load passengers was performed. The driver of the first minibus taxi was randomly selected from a list of taxi queue controllers, and then the driver of every third minibus taxi in the queue was requested to participate in the study. The population of drivers per taxi rank was approximately 110 and the total population was 990 drivers. A sample size calculated per taxi rank at 95% confidence level, 5% margin of error, and 50% response distribution [26] was 86 taxi drivers per rank. The total sample size reached for nine taxi ranks was 774 taxi drivers.

### Data collection

Data collection commenced from March to July 2016 after the Taxi Association and other key stakeholders from the different taxi ranks granted permission. Data collection was scheduled to occur during off-peak hours so that the research process did not interrupt the work of the taxi drivers. Data was collected by a team of trained fieldworkers using a researcher administered structured questionnaire translated from English into local languages. The drivers were asked about whether they would use patient-initiated PN, the likelihood of telling a partner that they themselves were diagnosed with STI, whether they would deliver a PN notification and referral slip to sexual partners, their acceptability of provider-initiated PN using SMS, and preference for notification if partner is diagnosed with STI. The tool further captures socio-demographic data, sexual behaviour, condom use, multiple sexual partnerships, and HIV testing history. The drivers were further asked a series of questions to assess the level of STI knowledge.

### Ethical considerations

The Sefako Makgatho Health Sciences University Research and Ethics Committee (SMUREC/H/284/2015: IR) granted ethical clearance. Permission to collect data was granted by the District Taxi association and the relevant management officials in destructive. Informed consent was obtained from drivers who volunteered to participate in the study prior to data collection. The taxi drivers were informed about voluntary participation and anonymity of their responses, personal information was not gathered from the drivers. Confidentiality and privacy were maintained during data collection. Four gazebos, each furnished with a table and two chairs were used to provide privacy while at the same time ensuring that data collection did not interrupt work in the taxi ranks.

### Data analysis

Descriptive statistics were conducted to describe demographics, sexual relationships, HIV testing practices, perceived use of referral slip from sexual partner, and level of acceptability of provider-initiated PN using SMS. Univariate and multivariate analyses using stepwise regression analysis were conducted to investigate association between dependant variables and independent variables.

Dependant variables were, perceived use of referral slip from sexual partner and acceptability of provider-initiated PN using SMS. The independent variables were demographic characteristics, sexual relationships, HIV testing history, condom use in the last sexual act, and likelihood of telling a partner when diagnosed with STIs. All the variables that showed significant association (p ≤ 0.05) at bivariate analysis were included in the stepwise logistic regression model. A p-value of less than 0.05 was considered significant. Results are expressed as odds ratios (ORs) with 95% confidence intervals (CIs) and p-values. All statistical analyses were performed using using STATA IC version 13.

## Results

A sample of 722 male minibus taxi drivers participated in the study and had been working in the minibus taxi industry for the mean period of 8.7 years (range, 1 to 40 years; SD=7.9 years). Their ages ranged from 19-68 years, with a mean age of 37.2 years (SD = 10.3 years). The majority of the minibus taxi drivers (43.8%) had received some secondary education but only 39.5% had completed secondary education (12^th^ grade) and only 8.6% has a tertiary education. Concerning living arrangements, 40.8% were married and only 36% lived with their wives in the same household while others left their wives and families and migrated for work in the City of Tshwane. On reports of sexual partners, 278 (44.2%) of the minibus taxi drivers had two sexual partners and 96 (15.3%) had three or more sexual partners. Concerning condom use, more than half (n=386, 52.2%) did not use a condom the last time they had sex and 117 (17.5%) reported never using a condom (Table 1).

**Table 1:**
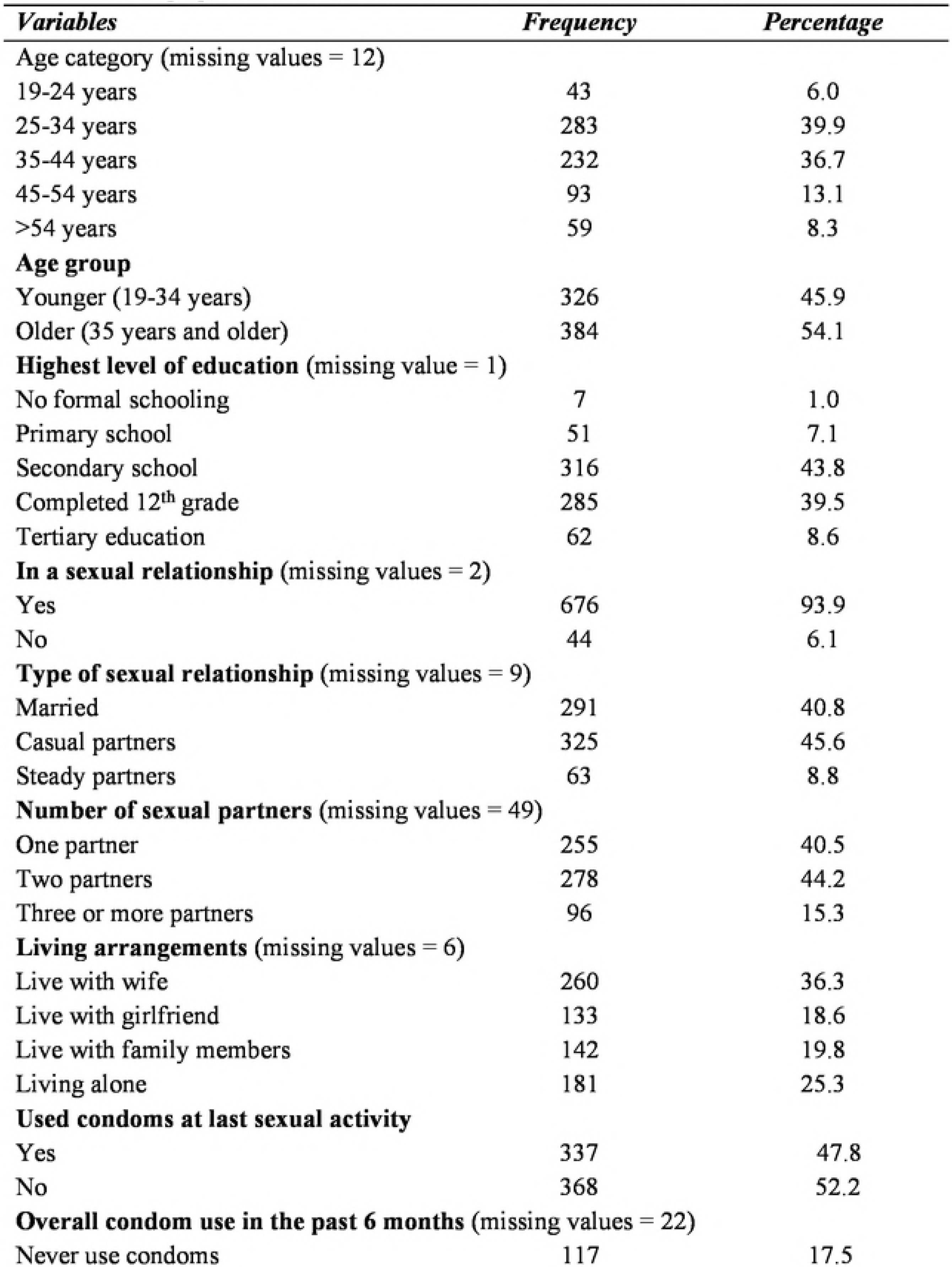
Demographic and sexual behaviour characteristics of minibustaxi driversin gauteng

| *Variables* | *Frequency* | *Percentage* |
| --- | --- | --- |
| **Age category** (missing values = 12) |  |  |
| 19-24 years | 43 | 6.0 |
| 25-34 years | 283 | 39.9 |
| 35-44 years | 232 | 36.7 |
| 45-54 years | 93 | 13.1 |
| >54 years | 59 | 8.3 |
| **Age group** |  |  |
| Younger (19-34 years) | 326 | 45.9 |
| Older (35 years and older) | 384 | 54.1 |
| **Highest level of education** (missing value = 1) |  |  |
| No formal schooling | 7 | 1.0 |
| Primary school | 51 | 7.1 |
| Secondary school | 316 | 43.8 |
| Completed 12<sup>th</sup> grade | 285 | 39.5 |
| Tertiary education | 62 | 8.6 |
| **In a sexual relationship** (missing values = 2) |  |  |
| Yes | 676 | 93.9 |
| No | 44 | 6.1 |
| **Type of sexual relationship** (missing values = 9) |  |  |
| Married | 291 | 40.8 |
| Casual partners | 325 | 45.6 |
| Steady partners | 63 | 8.8 |
| **Number of sexual partners** (missing values = 49) |  |  |
| One partner | 255 | 40.5 |
| Two partners | 278 | 44.2 |
| Three or more partners | 96 | 15.3 |
| **Living arrangements** (missing values = 6) |  |  |
| Live with wife | 260 | 36.3 |
| Live with girlfriend | 133 | 18.6 |
| Live with family members | 142 | 19.8 |
| Living alone | 181 | 25.3 |
| **Used condoms at last sexual activity** |  |  |
| Yes | 337 | 47.8 |
| No | 368 | 52.2 |
| **Overall condom use in the past 6 months** (missing values = 22) |  |  |
| Never use condoms | 117 | 17.5 |

### Knowledge of STI symptoms

Knowledge of STIs symptoms is a predictor for early access of treatment and partner notification and referral [21, 25). Almost all (n=692, 96.5%) of the minibus taxi drivers had awareness of STIs. For most (n=276, 39.6%), the source of STIs information was the clinic. Table 2 presents their responses about STI symptoms. Penile discharge as a most common STI symptom was reported by 270 (44.6%). The level of knowledge of STI symptoms was low, as only 319 (45.1%) knew that men could have STIs without symptoms at an early stage of infection. The results also showed that self-reported STI occurrence was low, with only 5.5% (n=40) of the participants reported to have been diagnosed with STIs in the past 12 months; this is despite the high occurrence of low usage of condoms among the study participants.

### Acceptability of STI partner notification

Table 3 presents the taxi drivers’ views about partner notification. Almost all (n=709, 98.2%) reported that it was important to notify a sexual partner about an STI diagnosis. A large proportion 698 (97.5%) were not sure whether they would verbally inform a sexual partner if they were diagnosed with STIs. In contrast, 670 (93.2%) would give a PN referral slip to a sexual partner if they were diagnosed with an STI, 504 (69.5%) felt that it would be easy to give a sexual partner a STI notification and referral slip and 659 (91.5%) said they would use a referral slip from their sexual partner to access treatment. The level of acceptability of PN using a SMS from a health care provider was 452 (62.7%). Over a third (n=234, 32.5%) found PN using SMS not acceptable, but only 109 gave reasons why PN using SMS is not acceptable. The reasons for non-acceptability of PN using SMS were, 55 (50.5%) drivers prefer telling their partner face to face, 47 (43.1%) said SMS is not reliable, and 7 (6.4%) said that SMS could cause conflict.

## Discussion

South Africa, as a developing country has opted for patient-initiated PN using referral slips as the current standard practices for STI control [11], while text messaging is deemed acceptable for provider-initiated PN in developed countries [13, 18]. The barriers to successful implementation of patient-initiated PN using referral slips were mentioned earlier in the manuscript. The concern is that South Africa has a high burden of STIs but it still relies on this strategy since the implementation of the syndromic management of STI in 2009 [27, 28].

Almost all of the minibus taxi drivers had awareness of STIs and accurately cited the STI symptoms even though more than 50% did not know that men could have STIs without showing symptoms during the early stages of the infection. The critical point of not knowing that STIs are asymptomatic during early stages of the infection explains the issue of missed opportunity in early infection, high STI transmission to sexual partners, and high prevalence of STIs in the country. This also has implications for the health education and health promotion programs as it shows there is a gap in creating awareness and accurate information on STI prevention, transmission, treatment, and control.

Risky sexual behaviour was determined by self-reports of non-condom use, with almost 20% of the minibus taxi drivers reporting to never have used condoms during sexual intercourse, over half did not use a condom during the last sexual act, and 42.8% reported inconsistent use of condoms. Moreover, two-thirds had more than one sexual partners and almost half (45.6%) were in casual sexual relationships. Inaccurate information and beliefs about STIs, having multiple sexual partners and risky sexual behavior were common in male STI patients as compared to female STI patients [12, 28, 29]. This argument also has implications for reviewing the content of health education and health promotion programs in order to close the knowledge gap among the high-risk groups.

A high proportion of the minibus taxi drivers knew the importance of notifying a sexual partner once diagnosed with STIs. However, there were conflicting views regarding informing sexual partners face-to-face and delivering a PN and referral slip should they be diagnosed with STIs. The majority (97.5%) were not kin to tell a partner face-to-face if they themselves were diagnosed with STIs but they prefer face-to-face delivery of the slip, which implies that they expect the sexual partner to take the slip without question or conversation. Lack of communication about sexual issues due to stigma and judgment was raised earlier in this manuscript and this has implications for STI treatment and control. This suggest that reproductive health programs such as STI treatment and control should explore more counselling and encourage STI couple counselling and treatment sessions.

There was a vast difference between perceived use of referral slip from a sexual partner using referral slip and the acceptability of PN using SMS from the HCWs (93% vs 62.7%). High interest in using notification and referral slips has shown to be less popular and less effective as a strategy to control STIs since only a small fraction of the slips were reported to be ever returned to clinics [30]. A systematic review of literature revealed that face-to-face patient-initiated PN interventions have had limited success which reflects discordance between high levels of acceptability of PN and low rates of utilization [14]. This is despite high level (70.2%) of PN counselling by HCWs and high acceptability and possible recommendation of PN to sexual partners [29]. There is evidence that both provider and patient-initiated PN informed less than 40% of sex partners [31]. The findings further suggest that PN could even be less in developing countries since the current PN and referral practices using referral slips fail to reach the majority of partners [16, 27, 28].

The relatively small difference between perceived easiness of delivering a PN referral slip to sexual partner, and acceptability of PN using SMS from a health care provider (69.5% vs 62.7%), with both being below 75% highlights the potential challenge of utilization of PN in this target group. Studies conducted in developing countries reported high levels of acceptability of PN with low rates of utilization and the findings of our study suggest the same. [19, 32]. This suggests that the minibus taxi drivers are not keen to disclose when they are infected and this implies that health educators need to do more work in promoting PN as a crucial intervention for prevention and control in this high-risk group.

A quarter of taxi drivers indicated personal preference of using SMS from health care providers over patient initiated PN referral slip. SMS, a form of electronic communication technology, has the potential to reduce costs, expand coverage, and increase efficiency of both provider– and patient-initiated PN services [18, 33, 34]. With some evidence of acceptability of provider-initiated PN using SMS, and where more than 3 out of 4 people in the world have mobile phones [18, 33], it would be beneficial as a compliment to PN slips in South Africa. With use of SMS, health care providers and patients can use mobile phones and text messaging services to contact partners in ways that were not possible prior to increased access with communication technologies [33, 34].

Evidence from studies indicate that the uptake of electronic PN has been slow, especially in settings where impact may be greatest [33]. In the current study, over a third minibus taxi drivers were not in favor of PN by SMS from a health care provider. They preferred telling partners face to face, they cited that SMS was not a reliable method and had a possibility of causing conflict in the relationship. These findings are in consensus with two other studies where face-to-face was one of the preferred method for contacting partners for STI treatment whereas SMS was less commonly used method [30, 33]. While electronic text messages are likely to increase notification to partners who may not be notified otherwise, research shows that individuals are more inclined to seek services when they are notified face-to-face rather than anonymously. It is important for health care providers to take note of the different preferences for PN to provide patients with more options to choose a PN method that is best suited to their relationships and circumstances [30, 34].

## Limitations

Our study findings should be interpreted taking the following limitations into consideration. The study sample consisted of a sample of taxi drivers in a district of one province in South Africa so the finding cannot be generalized to all minibus taxi drivers. This was a cross-sectional survey based on a hypothetical scenario assessing the perceptions of usage rather than actual usage and acceptability of partner notification as in randomized controlled trials. We could not ask for the reasons for the high acceptability of SMS provider initiated PN since this was a formative assessment of acceptability using a hypothetical scenario, therefore these findings do not provide an explanation of why the study sample told the partner of the STI diagnosis. Enhance comprehension of why the index case decide to inform or not inform the sexual partner about the STI will inform partner notification counselling messages provided in health facilities.

## Conclusion

We found that although high proportion of the minibus taxi drivers understood the importance of notifying a sexual partner once diagnosed with STIs, notifying the partner face to face was problematic. Consequently, delivering PN referral slip to sexual partner was preferable because it renders communicating about STIs and treatment the responsibility of the health care provider. The findings provide health care providers an opportunity to design partner notification STI messages to educate partners about STIs and encourage them to access treatment. Modifying the current patient-initiated PN will almost certainly serve the purpose of informing the partner about the STI with minimal involvement of the patient. Moreover, health facilities were the source of information about STI for all minibus taxi drivers, which suggest that health facilities are well positioned to increase access to PN and STI counselling.

Delivering a partner notification referral slip together with using a PN referral slip received from a sexual partner were preferred by the minibus taxi drivers. Another important finding is that over a third of minibus taxi drivers were not in favor of PN by SMS from a health care provider and preferred telling partners face to face. These findings raise concern on why now is the current patient initiated PN using referral slip is not working if it is the preferred method. This suggest that PN protocol should be flexible to allow health care providers to provide patients with options of a PN method that is best suited for the patient relationship and circumstance.

The challenges to notify the partner about a STI diagnosis face-to-face might explain the high acceptability of provider-initiated PN using SMS from a healthcare provider. Using SMS has an element of anonymity that delinks the index case from the STI diagnosis while offering the partner the opportunity to seek treatment. This would make sense in high-risk populations with multiple sexual partners and casual sexual relationships such as the current study sample.

With global expansion of mobile phone services, internet access, and reliance on the social media for finding sexual partners, it is essential that public health services develop strategies to increase utilization of both patient-initiated PN and referral using slips and provider initiated electronic notification systems in order to increase the numbers of partners notified to prevent the spread of STI significantly. In the context of South Africa where STIs continue to predispose individuals to HIV transmission, there is need for substantial efforts to support STI control interventions to ensure consistent implementation of patient-initiated PN.

## Abbreviations

HIV: Human Immunodeficiency Syndrome
PN: Partner Notification
SMS: Short Message Services
STI: Sexually Transmitted Infection

## Acknowledgements

This project would not have been possible without the willingness of the minibus taxi drivers who participated in this study, they really volunteered their time despite being in a busy competitive work environment. We thank the committed fieldwork team who gathered, managed and processed this data in preparation for analysis.

## Funding

This project received funding from the National Research Foundation (NRF) under the Grants application for non-rated scientists (CSUR150728131833) and VLIR/UOS, which is the Belgian University Consortium.

## Availability of data materials

A dataset will be submitted upon request by the Editor.

## Author contributions

MMM and SM conceived and designed the study, they were also responsible for the project implementation. MMM was responsible for data analysis and writing the manuscript. SM critically reviewed the manuscript and approved of it for publication.

## Competing interests

The authors declare that they have no competing interest.

## Consent for publication

The authors confirm that they have obtained consent to publish from the participants.

## Ethics

All participants were asked for written informed consent after the purpose of the study was explained to them. The research received ethical clearance from the Sefako Makgatho Health Sciences University Research and Ethics Committee (SMUREC/H/284/2015: IR).

